# An efficient experiment design helps to identify differential expressed genes (DEGs) in RNA-seq data in studies of plant qualitative traits

**DOI:** 10.1101/164202

**Authors:** Xingfeng Li, Zhenqiao Song, Yinguang Bao, Honggang Wang

**Affiliations:** State Key Laboratory of Crop Biology, Shandong Agricultural University, Taian, Shandong, China; Agronomy College, Shandong Agricultural University, Taian, Shandong, China

**Author notes:** Xingfeng Li and Zhenqiao Song contributed equally to the work. Corresponding author: Xingfeng Li and Honggang Wang.

**Keywords:** Comparative transcriptomics, qualitative traits, F_1_ hybrid, Dominant and recessive parent, Near-isogenic lines, *Triticum aestivum*

## Abstract

In this study, we conducted comparative transcriptome analysis between homozygous dominant parent and heterozygous F_1_ hybrid with homozygous recessive parent in qualitative trait study of common wheat (*Triticum aestivum* L.). Two sets of near-isogenic lines (NILs) were used: one set of NILs carrying powdery mildew resistance and susceptible *Pm2* alleles, the other set of NILs carrying different awn inhibition gene *B1* alleles. The results demonstrated that 2,932 DEGs were identified between L031 (*Pm2Pm2*) and Chancellor (*pm2pm2*), while 1,494 DEGs presented between F_1_ hybrid (*Pm2pm2*) and Chancellor, the co-regulated DEGs were 1,028. For the wheat awn inhibition gene *B1* test, 720 DEGs were identified between SN051-2 (*B1B1*) and SN051-1 (*b1b1*), and 231 DEGs were identified between F_1_ hybrid (*B1b1*) and SN051-1, the co-regulated DEGs were 180. Hierarchical clustering analysis of co-regulated DEGs showed that dominant parent and F_1_ hybrid were clustered as the nearest neighbors, while recessive parent showed an apparent departure. The results showed that the overlapping DEGs between dominant parent and F_1_ hybrid with recessive parent reduced the number of interested DEGs to only one-quarter (or one-third) of that between dominant and recessive parent, these overlapping loci could provide insights into molecular mechanisms that are affected by causal mutations.

## Introduction

RNA sequencing (RNA-seq) is an increasingly popular and effective approach for transcriptome analysis, with both high-throughput and high resolution capabilities for quantitative measurement of gene expression, detection of splice variants and single-nucleotide variations, and discovery of novel coding and noncoding transcripts (Shendure, 2008; Wang et al., 2009; Grabherr et al., 2011; Wang et al., 2014).

The identification of differentially expressed genes (DEGs) between two or more genotypes, tissues, time-points or conditions is a fundamental task in the analysis of RNA-seq data (Robles et al., 2012; Soneso and Delorenzi, 2013). The results obtained from the analysis of DEGs may facilitate our understanding of the molecular basis of phenotypic variation, disease resistance, developmental changes, and a diverse range of biological problems. Recently, some research groups have elucidated the performance characteristics of RNA-seq, different sources of measurement noise, and effects of different analysis approaches and algorithms (Kratz and Caminci, 2014; Li et al., 2014; Risso et al., 2014; Consortium 2014). Such studies have helped to improve the detection power of DEGs in RNA-seq analysis.

According to the theory of qualitative Mendelian traits, a cross between a homozygous dominant and a homozygous recessive line produces a heterozygous F_1_ hybrid that exhibits the same phenotype as the dominant parent. Therefore, the gene expression profiles of the causal genes and the affected downstream genes in the F_1_ hybrid would be similar to those of the dominant line. However, the F_1_ hybrid contains two alleles, one contributed by each parent; therefore, the differentially expressed genes between the F_1_ hybrid and the recessive parent could be very useful and more accurate for identifying important DEGs.

However, few studies have investigated experimental designs in this manner. Transcriptome studies of F_1_ hybrids and its parents have mostly been used to study phenomena associated with regulatory divergence (*cis/trans*), additive or transgressive gene expression, and allele-specific gene expression, thus providing an opportunity to speculate on mechanisms of heterosis (Paschold et al., 2012; Song et al., 2013; Bell et al., 2013; Song et al., 2014).

In this study, we conducted a comparative transcriptome analysis of the F_1_ hybrid with its parents for two qualitative traits of common wheat. These two RNA-seq experiments both revealed that DEGs between the F_1_ hybrid and recessive parent were clearly lower than that between the dominant and recessive parents, and the overlapping DEGs between them could provide more accurate information regarding the molecular mechanisms that are affected by the causal mutation.

## Materials and Methods

### Materials

Two sets of near-isogenic lines (NILs) of wheat were used: one set of NILs carrying powdery mildew resistance and the susceptible *Pm2* alleles, and the other set of NILs carrying differences in the wheat awn inhibition gene *B1*.

The wheat cultivar Chancellor (susceptible, *pm2pm2*) and its near-isogenic line L031 (resistant), which carries the powdery mildew resistance gene *Pm2*, were used in this study. They were obtained by crossing Ulka (donor of resistance gene) with Chancellor, and then backcrossed with Chancellor for 7 generations, and finally self-crossed and selected under *Blumeria graminis* f. sp. *tritici* inoculation (Wang et al. 2011). The plants of Chancellor, L031 and its F_1_ hybrid (resistant) were raised in a growth chamber with a 16-h photoperiod at 18°C. The *Bgt.* isolate E09 was used for infection at the trifoliate stage. Sixteen hours after inoculation, leaf samples were collected, pooled from three plants, immediately immersed in liquid nitrogen, and stored at −80 °C until processing. A pair of wheat lines with different awn lengths, SN051-1 (long awn, *b1b1*) and SN051-2 (short awn, *B1B1*), were used in the current study. They were developed from the F_8_ progeny of Octoploid *Triticale* Jinsong49 and Octoploid *Trititrigia* Xiaoyan7430, and then were self-crossed for 5 generations (Du et al. 2010). The morphological, genetic and molecular marker analyses showed that they differed in the wheat awn inhibition gene *B1*: line SN051-2 contained the dominant allele *B1*, and SN051-1 possessed the recessive allele *b1*. Genotypic data via the use of DNA markers showed that SN051-1and SN051-2 were truly isogenic (additional file 1). The hybrid was obtained from the cross of SN051-1 × SN051-2. The SN051-1, SN051-2, and F_1_ hybrid were planted in the same field at Shandong Agricultural University agronomy farm, Tai'an, Shandong, China. Pools of young spikes from ten plants were hand-dissected when they developed into the anther connective tissue formation phase, while the awn primordia at the distal end of the lemma began to elongate in the awned genotypes. The young spike samples of SN051-1, SN051-2, and the hybrid were collected, immediately frozen in liquid nitrogen, and then stored at −80 °C until use.

### MRNA isolation and library preparation for sequencing

Total RNA was extracted using the RNAsimple total RNA kit according to the manufacturer’s instructions (Tiangen, Beijing, China). The quantity and quality of the extracted RNA were verified by gel electrophoresis and spectrophotometry. Oligo(dT) - coated magnetic beads were used to enrich the mRNA, which was then broken into fragments using fragmentation buffer. Using these cleaved mRNA fragments as templates, first- and second-strand cDNAs were synthesized. The double-stranded cDNA was further purified using the QiaQuick PCR extraction kit (Qiagen, Hilden, Germany), resolved for final reparation and poly (A) addition, and then connected using different sequencing adapters. The expressed sequence tag (EST) libraries were constructed by PCR amplification after checking the quality by agarose gel electrophoresis and sequencing using an Illumina HiSeq™ 2000 platform by Biomarker Technology Co., Ltd (Beijing, China).

### *De novo* transcriptome assembly and annotation

To obtain clean reads, raw reads such as reads with adaptors, reads in which unknown bases represented more than 5% of the total bases, and low-quality reads (percentage of low-quality bases with a quality value □≤ □5 was more than 50% of a read) were removed. The clean reads were then assembled *de novo* into longer contigs based on their overlap regions using the Trinity platform. Different contigs from another transcript and their distance were further recognized by mapping clean reads back to the corresponding contigs based on their paired-end information, and thus, the sequences of the transcripts were determined. These were defined as unigenes. All of the unique sequences were annotated using BLASTX against the NCBI non-nucleotide (Nt) sequence database, NCBI non-redundant (Nr) protein database, Swiss-Prot, Gene Ontology (GO), Kyoto Encyclopedia of Genes and Genomes (KEGG), and Clusters of Orthologous Groups (COG) to predict and classify the functions.

### Annotation of expression

To evaluate the depth of coverage, all of the usable reads were realigned to each unitranscripts using Short Oligonucleotide Analysis Package (SOAP) aligner (http://soap.genomics.org.cn/soapaligner.html) and then normalized into RPKM values (reads per kb per million reads; Mortazavi et al., 2008). The DEG analysis was then performed using the Bioconductor package DESeq based on the ratio of the RPKM values. The false discovery rate (FDR) control method was used to identify the threshold of the P-value in multiple tests to compute the significance of differences in transcript abundance. In the present analysis, only unitranscripts with an absolute log_2_ ratio ≥ 1 and an FDR significance score < 0.01 were used for subsequent analyses. Hierarchical clustering and heat map generation were performed in R. The gene expression data were log_2_-transformed and then quantile-normalized prior to generating the heat map for direct comparison of the data.

### Validation of DEG expression by quantitative real time PCR (qRT-PCR)

Gene-specific primers were designed using Primer software (version 5.0) and the *Actin* gene was used as the internal control gene. The qRT-PCR reaction for target gene transcript amplification was carried out in a final volume of 25μL containing PCR buffer, 1 mM MgCl2, 0.2 mM dNTP, 1μL of SYBR green 1, 1U of Taq, 0.4 μM of each forward and reverse primers, and 2 μL diluted cDNA. The PCR reaction conditions were: denaturation at 95 °C for 5 min followed by 40 cycles of 95 °C for 10 s, annealing at the appropriate temperature (from 57 to 61 °C) for 30 s, extension at 72°C for 30 s, with a final extension at 72 °C for 10 min. All reactions were done in triplicate. The amplification data were analyzed using iQ5 software version 1.0 (Bio-RAD, USA). The threshold cycle (Ct) values of the triplicate PCRs were averaged and relative quantification of the transcript levels was undertaken using the comparative Ct method. The ΔCt value of the calibrator (the sample with the highest ΔCt value) was subtracted from every other sample to produce the ΔΔCt value, and 2^-ΔΔCt^ was taken as the relative expression level for each sample.

## Results

### RNA-seq results of wheat NILs with contrasting disease reactions

For the powdery mildew resistance gene *Pm2* test, leaf samples of the wheat cultivar Chancellor (susceptible, *pm2pm2*), the near-isogenic line L031 (resistance, *Pm2Pm2*) and its F_1_ hybrid were inoculated with *Bgt.* E09 and then used for RNA-seq analysis. After cleaning and checking the read quality, 4.16 Gb, 3.62 Gb and 4.06 Gb of clean data were generated, respectively. Among the clean reads, more than 90% had a read quality of Q30 (sequencing error rate, 0.1%) or higher. These reads were assembled *de novo* using Trinity platform software, resulting in 316,787 transcripts and 112,033 unigenes with an N50 length of 1,742 bp and 1,158 bp, respectively. Among these, 63,210 unigenes were annotated after BLAST searches using the Nr, Swiss-Prot, KEGG, COG and GO databases. The percentage of aligned reads mapping to unigenes in the library was generally approximately 80%, which indicated an acceptable quality of the aligned reads.

The gene expression levels were measured as reads per kilobase per million reads (RPKM), and putative DEGs were identified using a false discovery rate (FDR) of less than 0.01 and a fold change greater than 2. The box plots of the relative log RPKM values for each RNA-seq library showed little distributional differences among the three libraries (Fig. 1A). As shown in Figure 1B, comparison of the L031 and Chancellor libraries (Pm vs. Ch) showed that 2,932 genes (account for 4.64% of annotated unigenes) were differentially expressed, 1,438 and 1,494 of these DEGs were up- and down-regulated after inoculation, respectively. Concomitantly, 1,419 DEGs (account for 2.24% of annotated unigenes) were identified from ‘F_1_ vs. Ch’ comparison, 692 DEGs were up-regulated, and 727 DEGs were down-regulated, respectively. Venn diagram of the differentially co-regulated genes revealed a total of 1,028 DEGs (account for 1.63% of annotated unigenes) presented in comparison between resistant (L031 and the F1) and susceptible genotypes (Chancellor). Four hundred and seventy-seven DEGs showed relatively higher transcript levels in resistant seedlings than susceptible seedlings, while 551 DEGs showed the opposite expression pattern (Fig. 1B).

**Figure 1.**
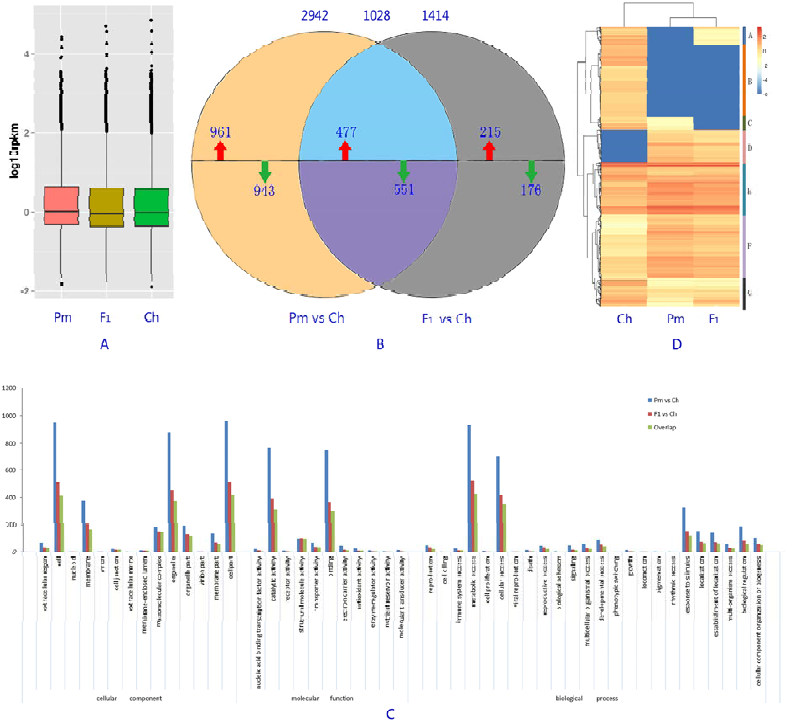
RNA-seq analysis of L031, Chancellor and its F_1_ hybrid after inoculation with ***Bgt.*** A. Box plots of the relative log FPKM values based on all of the genes for each RNA-seq library. B.Venn diagram showing the number of differentially co-expressed genes among Chancellor (susceptible, *pm2*), L031 (resistant, *Pm2*) and its F_1_ hybrid. (Red and blue arrows indicate up-and down-regulated DEGs, respectively). C.GO classifications of DEGs. The results are summarized into three main categories: ‘biological processes, cellular components, and molecular functions’. D.Heat map showing all of the DEGs among the three samples. The gene expression data were log_2_-transformed and then quantile-normalized prior to generating the heat map for direct comparison of the data. DEGs (red and blue indicate up-and down-regulated DEGs, respectively) for each sample were mapped by lane. Hierarchical clustering and heat map generation were performed in R.

GO enrichment analysis were conducted in order to determine the functions of the DEGs, these DEGs were categorized into 49 functional groups in the three main categories, “cellular components (CC)”, “molecular functions (MF)” and “biological processes (BP)” (Figure 1C). The term “cell“, “organelle” and “cell part” were significantly enriched in the CC category, term “catalytic activity” and “binding” were significantly enriched in the MF category, term “metabolic process” and “cellular process” were significantly enriched in the BP category. GO analysis also showed DEG numbers enriched in most terms predicted between F_1_ and Chancellor were reduced to only 1/2 or 2/3 of that between L031 and Chancellor, while term “macromolecular complex” in CC and term “structural molecule activity” did not (Fig. 1C).

Hierarchical clustering analysis of 1028 co-regulated DEGs in Chancellor, L031 and the F_1_ hybrid was performed (Fig. 1D). L031 and F_1_ hybrid were clustered as the nearest neighbors as anticipated, while Chancellor showed an apparent departure. Cluster analysis of the RPKM data showed that the differentially co-expressed DEGs could be grouped into seven clusters; clusters D and F were enriched with transcripts that showed large fold changes in expression between resistant and recessive genotypes.

Eight transcripts, which encoding the LRR family of receptor serine/threonine-protein kinases and up-regulated by *Bgt.* infection in the resistant genotypes (L031and F_1_), were selected for quantitative real-time PCR (qRT-PCR) analyses (Fig. 2). The RPKM values of these transcripts were much lower (some were 0) in Chancellor, hence these genes may be responsible for perception of fungal invasion and activation of down-steam signaling cascades to induce defense responses. qRT-PCR analyses showed that the expression patterns of seven genes were highly consistent with the RNA-Seq data, suggesting that our transcriptome analysis was accurate and reliable.

**Figure 2.**
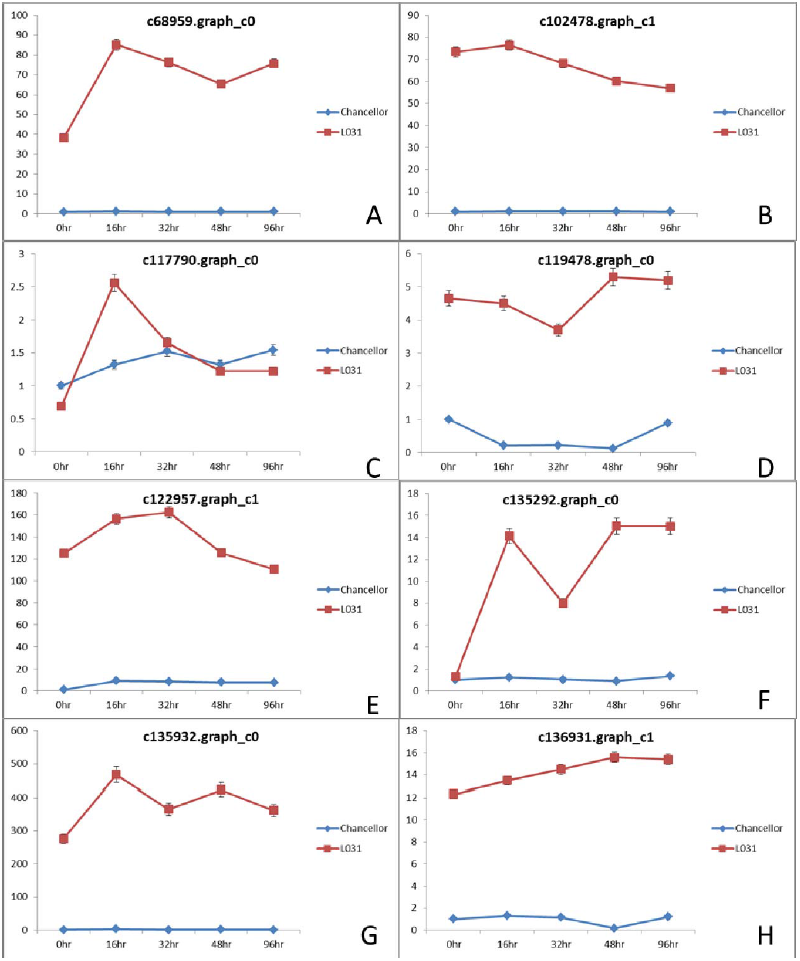
Validation of 8 DEGs in L031 and Chancellor using quantitative real time RT-PCR (qRT-PCR). The x-axis shows the samples at different hours after inoculation with *Bgt.* E09.

### RNA-seq results of wheat NILs that differ in wheat awn inhibition gene ***B1***

For the wheat awn inhibition gene *B1* test, global transcriptional changes of the young spike samples at the anther connective tissue formation phase of wheat lines SN051-1 (awned, *b1b1*) and SN051-2 (awnless, *B1B1*), and its hybrid were compared. Approximately 9.34, 9.18 and 10.27 Gbs of clean data were generated using the Illumina HiSeq™ 2500 sequencing device, respectively. The Q30 values were higher than 85.0%. *De novo* assemblies using the Trinity program generated 265,244 transcripts and 96,360 unigenes with an N50 length of 1,628 and 1,231 bp, respectively. A total of 49,289 unigenes were annotated.

Transcriptome profiles of young spike samples of SN051-2 (Al), SN051-1 (An) and its F_1_ hybrid were compared in order to obtain a better understanding of awn development at the transcriptome level. The box plots of the relative log FPKM values based on all of the genes showed profiles similar to the profiles obtained in the *Pm2* test (Fig. 3A). A total of 720 and 231 DEGs were identified by pairwise comparisons of the RNA-seq data between ‘SN051-2 vs. SN051-1’ (Al vs. An) and ‘F_1_ hybrid vs. SN051-1’ (F_1_ vs. An), with 549 and 168 up-regulated and 171 and 63 down-regulated genes, respectively (Fig. 3B). The overlap analysis between ‘Al vs. An’ and ‘F_1_ vs. An’ identified 180 DEGs (account for 0.36% of annotated unigenes) that showed constant differential expression (134 up-and 46 down-regulated) (Fig. 3B), representing only one-quarter of the DEGs between dominant and recessive parent.

**Fig. 3.**
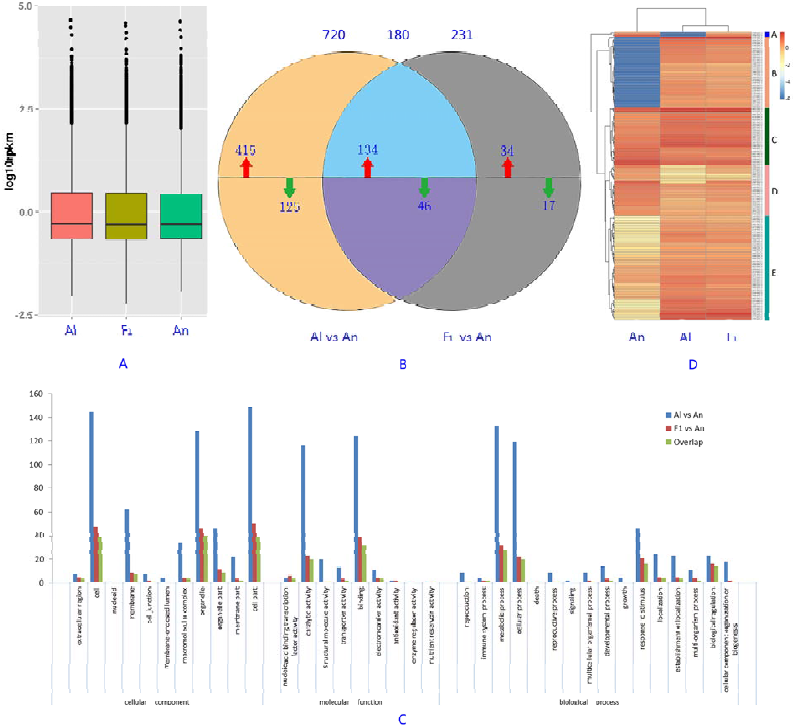
RNA-seq analysis of young spikes of SN051-2 (Al), SN051-1 (An) and its F_1_ hybrid. A. Box plots of the relative log FPKM values based on all of the genes in each RNA-seq library. B. Venn diagram showing the number of differentially co-expressed genes among Chancellor (susceptible, *pm2*), L031 (resistant, *Pm2*) and its F_1_ hybrid. (Red and blue arrows indicate up-and down-regulated DEGs, respectively). C. GO classifications of DEGs. The results are also summarized into three main categories: ‘biological processes, cellular components, and molecular functions’. D. Heat map showing all of the DEGs among three samples. The gene expression data were log_2_-transformed and then quantile-normalized prior to generating the heat map for direct comparison of the data. DEGs (red and blue indicate up-and down-regulated DEGs, respectively) for each sample were mapped by lane. Hierarchical clustering and heat map generation were performed in R.

GO enrichment analysis revealed that interested DEGs between the awnless and awned genotypes were categorized into 36 functional groups in CC, MF and BP categories. The term “cell”, “organelle” and “cell part” in the CC category, term “catalytic activity” and “binding” in the MF category, and term “metabolic process” and “cellular process” in the BP category were also significantly enriched (Figure 3C). DEG numbers between F_1_ and SN051-1 enriched in most terms were significantly down-regulated than that between SN051-2 and SN051-1.

Hierarchical cluster analysis showed that dominant parent SN051-2 and F_1_ hybrid tended to cluster together as expected, which revealed similar overall gene expression profiles between them (Fig. 3D). Clustering analysis arranged 180 significantly DEGs into five groups. Cluster B was enriched with 44 transcripts that were uniquely expressed in the awnless genotype, and although most of the results were hypothetical proteins, they might be considered genes involved in regulation of awn development. However, this hypothesis requires further investigation.

Among 46 down-regulated DEGs (overlapped between ‘Al vs. An’ and ‘F_1_ vs. An’), four transcripts were selected for quantitative real-time PCR (qRT-PCR) analyses (Fig. 4). Results showed gene expression trend of these DEGs were also highly consistent with to RNA-Seq data.

**Figure 4.**
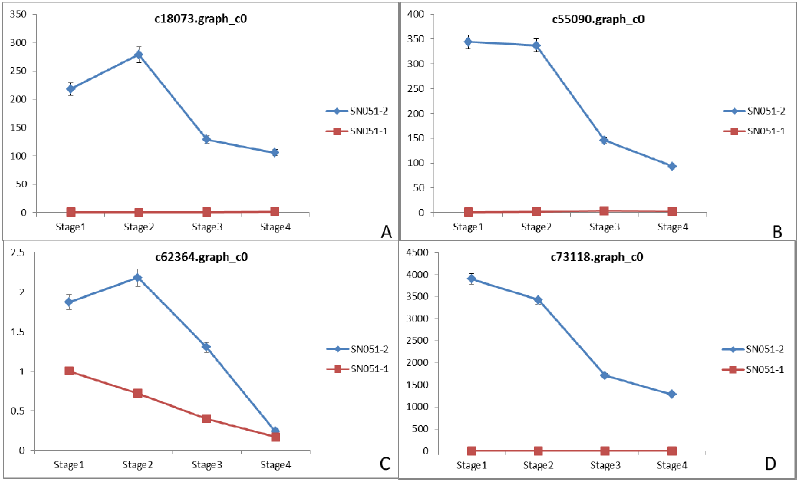
Validation of 4 DEGs in SN051-1 and SN051-2 using quantitative real time RT-PCR (qRT-PCR). The x-axis shows the samples at different develop stage.

## Discussion

Although differences in expression between wild-type and mutant isolates can help to answer biological questions, DEG analysis does not generally facilitate the identification of an initial list of potential candidate genes. Even in RNA-seq and DEG analyses of qualitative traits, thousands of DEGs have been identified between near-isogenic lines (Zhou et al., 2010; Zhang et al., 2014). Some of the DEGs are likely from the result of measurement noise or biological complexity, whereas a wide range of the DEGs likely demonstrate the downstream effects of the causal genes (Cai et al., 2014; Lin et al., 2014). An increase in the sequencing depth and numbers of samples would decrease these biases to some extent (McIntyre et al., 2011; Rapaport et al., 2013).

In the qualitative Mendelian traits study, F_1_ hybrid exhibits the same phenotype as the dominant parent, the gene expression profiles of the causal genes and the affected downstream genes between the F_1_ hybrid and its dominant parent would be similar. In that means, the DEGs presented between the F_1_ hybrid and the recessive parent could be very useful and more accurate for identifying important DEGs. In this study, according to all of the DEGs statistics for the 4 pairwise comparisons in the two treatments, we found that the DEG numbers between the F_1_ and recessive parent was clearly less than that between the dominant and recessive parent. Using the DEGs that overlapped between the homozygous dominant parent and heterozygous F_1_ hybrid with the homozygous recessive parent condensed the interested DEGs to 1/3 or 1/4 of DEGs between the dominant and recessive parent.

This finding indicated that we could compare the F_1_ generation with the susceptible parent to reduce the number of false positive genes and improve the veracity of the experiment. In other words, although this approach does not provide a direct identification of the 'Trigger' gene of interest, it brings us closer to the answer.

Previous studies proved that differential gene expression in F_1_ hybrid and its parents may be very complex. For example, F_1_ hybrid could accumulate levels of transcript equal to the mid-parent (additivity), the high or low parent (high or low parent dominance), above the high parent (over-dominance), or below the low parent (under-dominance) (Swanson-Wagner et al., 2006). Recent studies have demonstrated that mono-allele specific expression play important roles in development and stress induced responses, thus contributed to heterosis in plants (Song et al., 2013; Song et al., 2014; Guo et al., 2015). The object of the present study were to study the molecular mechanism of the qualitative traits, but we should not ignored the fact that differential expressed genes between F_1_ hybrid and parents could be the results of induction and suppression of the genes caused by genetic and epigenetic alteration. These genes also possess the biological information, some of these DEGs should be studied in detail while should not be considered as the false positives in hasty.

## Conclusions

Two different RNA-seq experiment of qualitative trait in wheat showed that the differentially expressed genes between F_1_ and recessive parent were lower than those observed between dominant and recessive parents, overlapping loci between the homozygous dominant parent and heterozygous F_1_ hybrid with the homozygous recessive parent could reduce the number of interest DEGs to only one-quarter (or one-third) of the total number of DEGs between dominant and recessive parent, which would provide more accurate information regarding the molecular mechanisms that are affected by the causal mutation.

## Additional file

Additional Table S1. Percentage of the polymorphic markers between L031 and Chancellor. Additional Table S2. Percentage of the polymorphic markers between SN051-1 and SN051-2. Additional Table S3. The 1,028 overlap DEGs in the comparison between resistant (L031 and the F1) and susceptible genotypes (Chancellor) in *Pm2* test.

Additional Table S4: The 180 overlap DEGs between ‘Al vs. An’ and ‘F_1_ vs. An’.

Additional Figure S1. Expression patterns of the 8 R protein-like transcripts which were up-regulated in L031 and F1 by RNA-seq data

Additional figure S2: Expression patterns of the 4 DEGs which were down-regulated in SN051-1 by RNA-seq data.

## Acknowledgments

This work was supported by The National Key Research and Development Program of China (2016YFD0102000) and National Natural Science Foundation of China (31671675), Natural Science Foundation of Shandong Province (ZR2015CM034).

